## Supplementary figures and images for "From Movement to METs: A Validation of ActTrust® for Energy Expenditure Estimation and Physical Activity Classification in Young Adults"

### Supplementary Figure S1

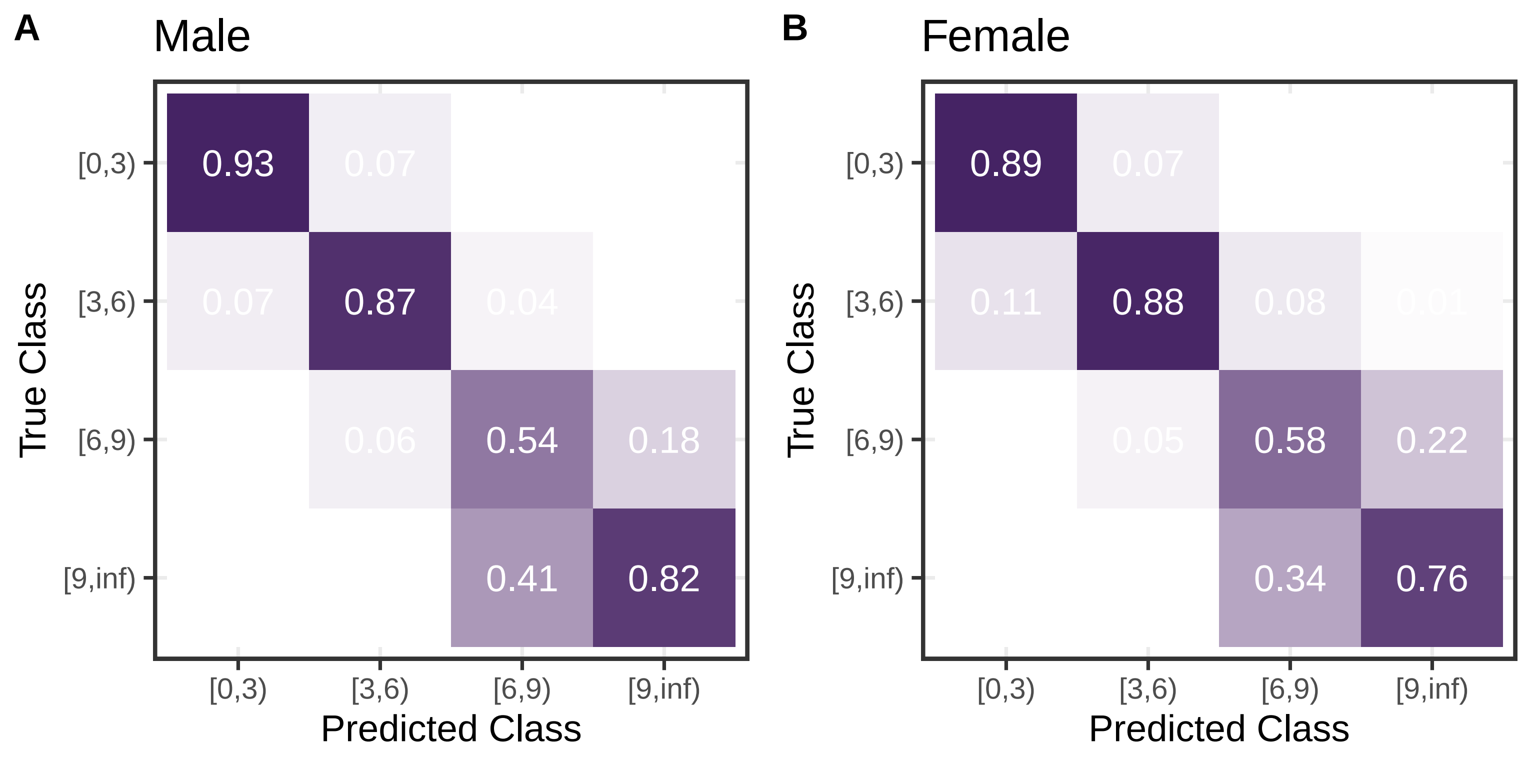

### Supplementary Figure S2

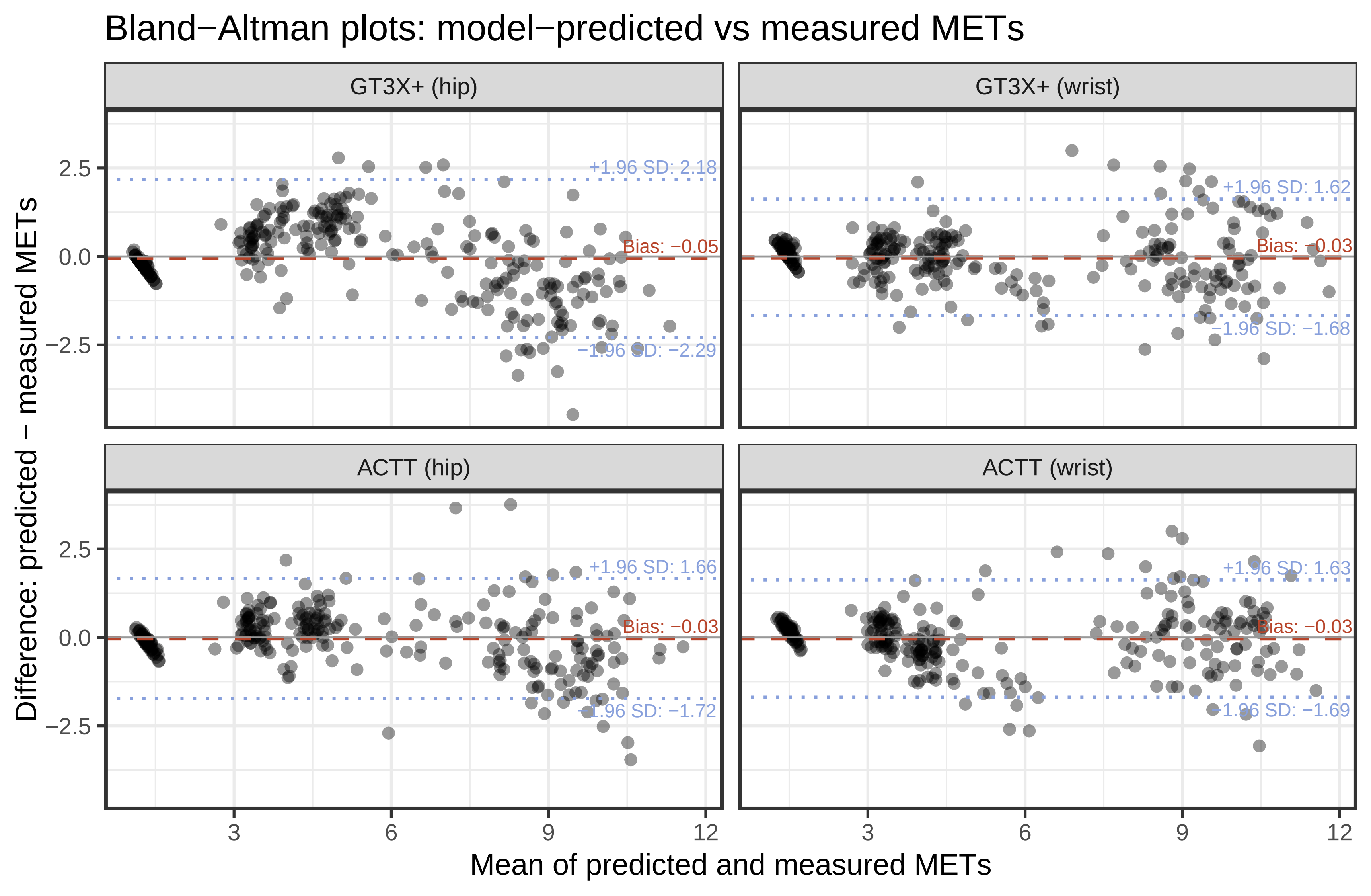

### Supplementary Figure S3

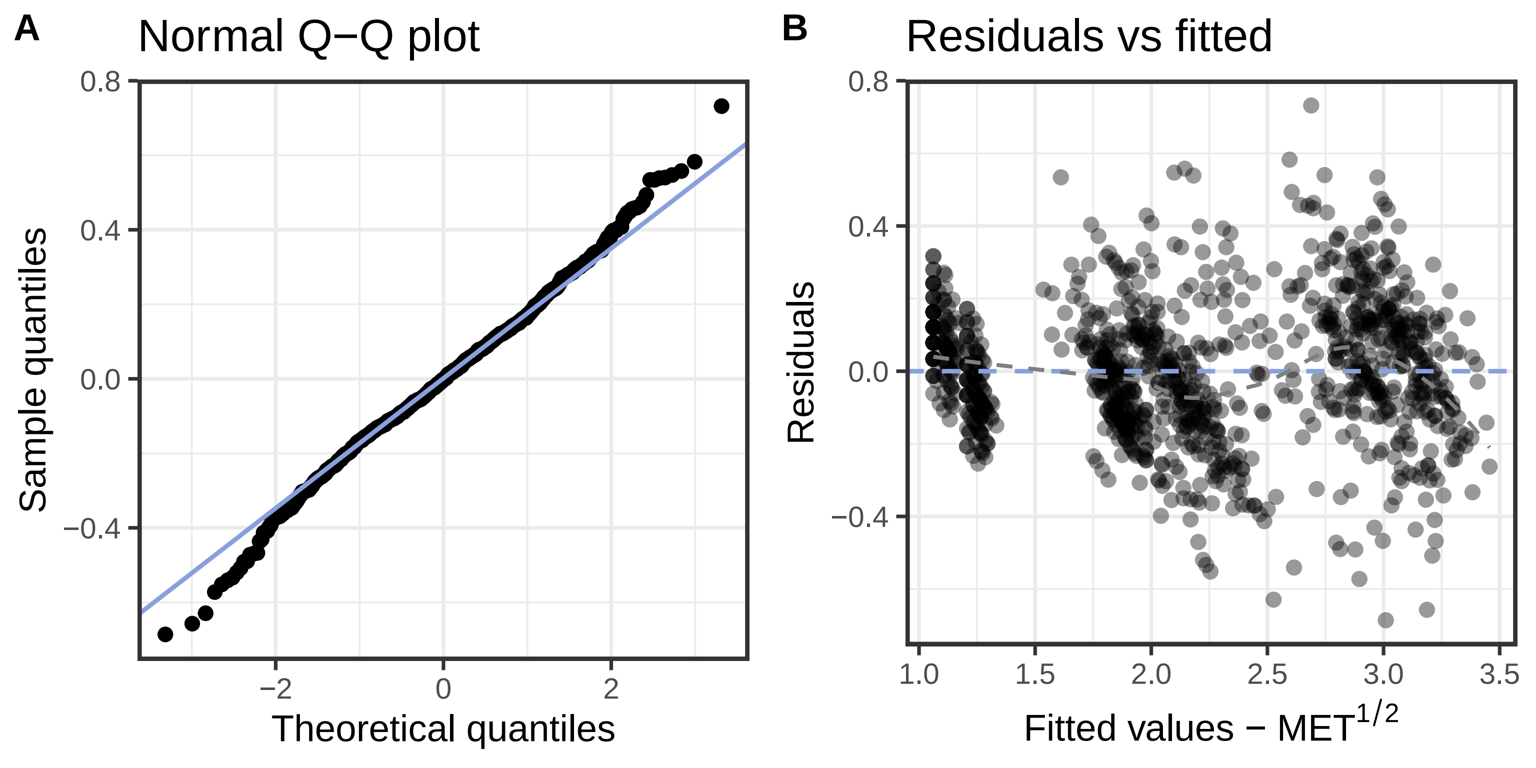
